# Bioinformatic analysis of the spike protein cleavage sites of coronaviruses in the mammalian order Eulipotyphla

**DOI:** 10.1101/2023.02.05.527216

**Authors:** Qinghua Guo, Annette Choi, Jean K. Millet, Gary R. Whittaker

## Abstract

The mammalian order Eulipotyphla, including hedgehogs and shrews, represent a poorly understood reservoir of coronaviruses with zoonotic potential. Here, we carried out a bioinformatic analyses of these viruses based on the viral spike protein—to illustrate the complexity of coronavirus evolutionary history and the diversity of viruses from these host species, with a focus on the presence of possible furin cleavage sites within the spike protein. We found no evidence for cleavage by furin itself; however, certain strains of Wencheng Sm Shrew coronavirus were shown to have a predicted cleavage site for other member of the proprotein convertases, which are furin family members— suggesting their spillover potential. As the expanding urbanization and the trade of small mammals in the wet markets enhance the wildlife-human interactions, this may increase pathogen spillover risks. Therefore, we should implement broad wild animal surveillance and be vigilant of contact with these small wild mammals in light of one-health perspectives.

## Introduction

The current global pandemic of severe acute respiratory syndrome coronavirus 2 (SARS-CoV-2), named COVID-19 by WHO, is having a huge impact on global health (Rabaan et al., 2020). It emerged 7 years after an outbreak of Middle East respiratory syndrome coronavirus (MERS-CoV) and 16 years after an outbreak of severe acute respiratory syndrome coronavirus (SARS-CoV) (van Boheemen et al., 2012). The Coronaviridae Study Group (CSG) of the International Committee on Taxonomy of Viruses officially classified SARS-CoV-2 within the family *Coronaviridae* (“ The Species Severe Acute Respiratory Syndrome-Related Coronavirus,” 2020).

Coronaviruses are enveloped positive-strand RNA viruses and members of the subfamily *Orthocoronavirinae* in the family *Coronaviridae* within the order *Nidovirales* (Brian & Baric, 2005). Coronaviruses are classified into four genera, *Alphacoronavirus, Betacoronavirus*, and *Gammacoronavirus*, and *Deltacoronavirus*. Overall, there are currently seven well-recognized human coronaviruses, HCoV-229E, HCoV-NL63, HCoV-OC43, HCoV-HKU1, MERS-CoV, SARS-CoV, and the most recent SARS-CoV-2 (Decaro & Lorusso, 2020). HCoV-229E and HCoV-NL63 belong to the genus *Alphacoronavirus* while all the rest are classified as *Betacoronavirus*. Specifically, *Betacoronavirus* is further divided into four monophyletic lineages A-D (van Boheemen et al., 2012). Lineage A (embecovirus) includes HCoV-OC43 and HCoV-HKU1; lineage B (sarbecovirus) includes SARS-CoV and SARS-CoV-2 ; MERS-CoV and related viruses belong to lineage C (merbecovirus).

Interestingly, all the viral strains in lineage A and B are detected in different host species while those in lineage C and D are found in bats. For instance, HCoV-OC43 was transmitted to humans through domestic animals such as cattle or pigs (Corman et al., 2018); HCoV-229E derived from bats is hosted by alpacas (Corman et al., 2015); the ancestors of HCoV-NL63 are circulating in bats, whereas HCoV-HKU1 originated in rodents (Tao et al., 2017). SARS-CoV and MERS-CoV are also originated in bats and are carried by a wide range of wild animals (Decaro & Lorusso, 2020).

The size of coronavirus genome RNA is among the largest known so far. Despite the presence of a proof-reading EndoU enzyme, the diverse genetic features of coronaviruses are due to a high level of recombination and mutation, facilitating the emergence of novel viruses. Horseshoe bats are considered the most likely natural reservoir for SARS-CoV-2. However, the intermediate hosts for SARS-CoV-2 have still not been definitively identified, and these hosts could be a range of highly diversified animals, including mammals, avians, and reptiles (Boni et al., 2020; Hedman et al., 2021). Both domestic animals, such as dogs, cats, and pigs, and wildlife, such as pangolins, mink, and ferrets, can be intermediate hosts for SARS-CoV-2 (Zhao et al., 2020). Therefore, it is extremely important to continue studying coronavirus in wild small mammals and their zoonotic potential.

As mentioned above, the insectivorous bats under the order Chiroptera are important hosts for these alphacoronaviruses and betacoronaviruses. The ability to mutate and recombine enables coronaviruses to jump through different animal species (Cui et al., 2019). Members of the mammalian taxa of the order Eulipotyphla, including the hedgehogs, shrews, moles, and solenodons, have been demonstrated to have a close genetic relationship with the order Chiroptera (Tsagkogeorga et al., 2013) and are also insectiviorous. Since the first identification of coronaviruses in European hedgehogs (*Erinaceus europaeus*) in German, EriCoV (Erinaceus Coronavirus) was detected in France, the United Kingdom, and Italy (Corman et al., 2014; De Sabato et al., 2020; Monchatre-Leroy et al., 2017). Furthermore, another hedgehog coronavirus was found in Amur hedgehogs (*Erinaceus amurensis*) from China (Lau et al., 2019). Several Wencheng Sm shrew coronaviruses (WESVs) were isolated in the Asian house shrews (*Suncus murinus*) while Shrew coronavirus was discovered in the common shrews (*Sorex araneus*) in China (Wang et al., 2017; Wu et al., 2018).

The Rodentia order is believed to contain the most zoonotic host species, followed by Chiroptera and then two families within the order Eulipotyphla, the shrews (Soricidae) and the moles (Talpidae) (Han et al., 2016). However, there are far fewer studies focusing on the Eulipotyphla mammals compared to the other insectivores like bats, which might cause the gap in understanding these small mammal host species. These small mammals, always living in a complex and densely populated community with rich species, also share similar biological characteristics of high metabolic rates (Bray et al., 2008; Han et al., 2016).

The coronavirus spike (S) glycoprotein on the viral surface plays a vital role in viral infection (Millet et al., 2016; Wang et al., 2013). For instance, it has been demonstrated that MERS-CoV spike protein can adapt to the DPP4 (the cellular receptor for MERS-CoV) variation of different hosts by altering its surface charge (Letko et al., 2018; Wang et al., 2013). The spike protein is composed of the S1 domain in the N-terminal region and is followed by the S2 domain (Wang et al., 2013). The S1 domain helps the viral entry into the target cell by binding to the host cell receptor; the S2 domain mediates membrane fusion (Belouzard et al., 2009). Overall, the spike protein can be proteolytically cleaved and activated at two cleavage sites (Millet & Whittaker, 2014). The cleavage occurs at the boundary between S1 and S2 (S1/S2) during biosynthesis of the spike while at the upstream position of the fusion peptide (S2’) during viral entry. The N-terminal domain (NTD) and C-terminal domain (CTD) in S1 are likely to serve as the receptor-binding domain (RBD) (Li, 2016). The majority of S1-NTDs bind carbohydrates, except in MHV, which can recognize the receptor CEACAM1. S1-CTDs are responsible for binding receptors, including ACE2, APN, and DPP4, via the RBD (Wang et al., 2013). As enveloped viruses, these coronaviruses enter host cells by membrane fusion. Therefore, the proteolytic cleavage and the proteases cleaving and activating the spike protein play significant roles in understanding the viral pathogenesis. Although cellular entry might be different between Eulipotyphla coronaviruses and MERS-CoV/SARS-CoV/SARS-CoV-2 based on their low similarities in the corresponding RBD, it is still worthy to study the zoonotic potential of Eulipotyphla coronaviruses. This research might provide more insights into the viral transmission between the wild small mammals including bats, hedgehogs, and shrews, as well as the evolutionary origins of coronaviruses. We also want to highlight the importance of the wild small mammals as zoonotic host species, especially these insectivores from the order Eulipotyphla.

Here, we studied the phylogenetic relationships within Eulipotyphla coronaviruses and related coronaviruses. We also predicted the cleavage sites in the Eulipotyphla coronaviruses and explored the relationships of furin across different animal species. It was our goal to better understand the zoonotic potential of these wildlife-related coronaviruses and provide some suggestions for the public health field to prevent viral zoonoses.

## Methods

### Phylogenetic Analysis

The multiple sequence alignment was constructed in Geneious Prime 2020. The maximum likelihood phylogenetic tree was built in Mega-X 10.2.4 (Kumar et al., 2018). All viral spike protein amino acid sequences were downloaded from the NCBI protein database (https://www.ncbi.nlm.nih.gov/protein/). The spike protein accession numbers on the NCBI database were: Shrew-CoV/Tibet2014 (YP_009755839.1); Wencheng Sm shrew coronavirus isolates: Yudu-76 (ASF90460.1), Yudu-19 (ASF90465.1), Ruian-90 (ASF90470.1), Wencheng-554 (ASF90486.1), Wencheng-578 (ASF90491.1), Wencheng-562 (ASF90496.1), Xingguo-101 (YP_009389425.1), Xingguo-74 (YP_009824974.1); Rhinolophus bat coronavirus HKU2 (ABQ57208); Swine acute diarrhea syndrome coronavirus (AXY04083); Erinaceus hedgehog coronavirus cw_6c (QOQ34381); Betacoronavirus Erinaceus isolates: Italy/116988-1/2018 (QRN68024), Italy/50265-19/2018 (QRN68031), Italy/50265-17/2018 (QRN68055), Italy/50265-1/2018 (QRN68048), Italy/50265-11/2019 (QRN68066), Italy/50265-12/2019 (QRN68078), Italy/50265-13/2019 (QRN68090), Italy/50265-15/2019 (QRN68101), VMC/DEU/2012 (YP_009513010.1); Hedgehog coronavirus 1 (QCC20713.1); Erinaceus hedgehog coronavirus HKU31 (QGA70702.1); Betacoronavirus England 1 (K9N5Q8); BtVs-BetaCoV/SC2013 (AHY61337.1); Bat coronavirus HKU5-1 (ABN10875.1); Bat coronavirus HKU4-1 (ABN10839.1); Hypsugo bat coronavirus HKU25 (ASL68953.1); Pipistrellus abramus bat coronavirus HKU5-related (QHA24687.1); Tylonycteris pachypus bat coronavirus HKU4-related (QHA24678.1).

### Geographical Mapping of Coronaviruses from the Mammalian Order Eulipotyphla

The maps were spatialized using QGIS Desktop 3.16.3. The location information of the shrew coronavirus and hedgehog coronavirus isolates included in the map was retrieved from NCBI viral genome database (https://www.ncbi.nlm.nih.gov/).

### Furin Cleavage Sites Prediction

Furin cleavage site prediction was generated by the ProP 1.0 Server (https://services.healthtech.dtu.dk/service.php?ProP-1.0).

### Spike Protein Model Construction

The spike protein models were built using Rosetta Comparative Modeling (Song et al., 2013). The reference spike protein structure from the coronavirus HKU2 (PDB: 6M15) was downloaded from RCSB Protein Data Bank (https://www.rcsb.org/).

### Mammalian Furin Sequence Alignment

The multiple sequence alignment of the mammalian furin was performed by ClustalW (https://www.genomse.jp/tools-bin/clustalw) (Thompson et al., 1994).

The mammalian furin amino acid sequences were obtained from NCBI protein database (https://www.ncbi.nlm.nih.gov/gene/5045/ortholog/?scope=7776). The protein Accession numbers on NCBI database were: rat (NP_001074923.1); dog (XP_850069.2); pig (XP_001929382.1); cat (XP_023110662.1); giant panda (XP_034523748.1); rabbit (XP_002721548.2); camel (XP_031296453.1); human (NP_002560.1); cattle (XP_024837365.1); horse (XP_005602832.1); ferret (XP_004763758.1); fruit bat (XP_036087816.1); greater horseshoe bat (XP_032955811.1); dolphin (XP_004322334.1); shrew (XP_004617631.1); hedgehog (XP_016043802.1)

## Results and Discussion

The Eulipotyphlan relationships are depicted in Figure 1.

**Figure 1.**
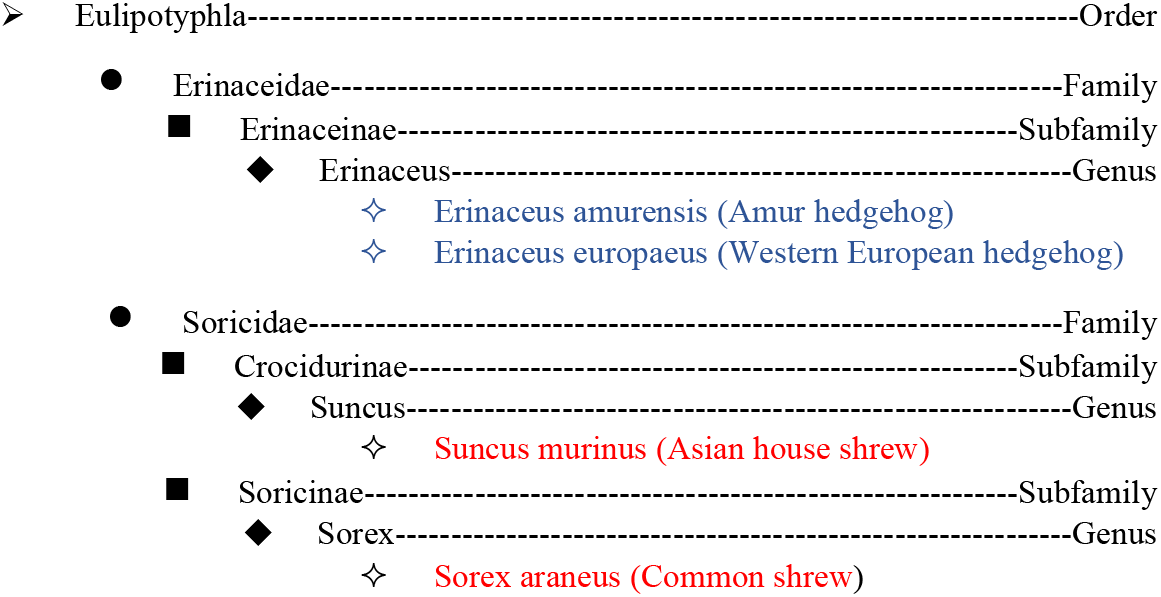
Eulipotyphlan relationships including coronavirus host species. The hedgehog species are colored in blue, and the shrew species are colored in red.

### Eulipotyphlan Coronavirus Host Species Relationships

Coronaviruses have been identified in hedgehogs and shrews in the order Eulipotyphla. The two hedgehog species are both classified in the genus Erinaceus belonging to the subfamily Erinaceinae under the family Erinaceidae. Western European hedgehogs included in the study were from Germany, France, Italy, and the UK (Corman et al., 2014; De Sabato et al., 2020; Monchatre-Leroy et al., 2017). Amur hedgehog (Erinaceus amurensis) coronaviruses were only detected in Jiangsu Province and Guangdong Province in China (Lau et al., 2019). The two shrew species both belong to the family Soricidae but in different subfamilies. Asian house shrews (*Suncus murinus*) are in the genus Suncus in the subfamily Crocidurinae, whose coronaviruses were discovered in Jiangxi Province and Zhejiang Province in China (Wang et al., 2017). The common shrews are within the genus Sorex and the subfamily Soricinae, whose coronaviruses were only detected in Tibet Autonomous Region in China (Wu et al., 2018).

### Phylogenetic Analysis of the Spike Protein Amino Acid Sequences

To understand the phylogenetic relationship between Eulipotyphla coronaviruses and other related coronaviruses, we constructed a maximum likelihood phylogenetic tree based on the spike amino acid sequences. In doing this, we also wanted to classify those strains with uncertain categories.

The classification represented in Figure 2 was obtained from the current NCBI taxonomy browser under the subfamily Orthocoronavirinae. All EriCoVs within the same species *Erinaceus europaeus* found in Italy, Germany, and the UK share an 89.96% to 100% spike amino acid sequence identity. All of the unclassified Italian EriCoVs are similar to the other Eri-CoV isolates from Germany and the UK with 100% bootstrap support. Therefore, these Italian EriCoVs are likely to be classified as Merbecovirus. The two Amur hedgehog coronavirus HKU31 strains from China form a single clade distinct from the cluster of EriCoVs with 77.61% to 79.1% spike amino acid sequence identity. EriCoV and HKU31 spike proteins have an average identity of 55.84% and 56.6% with MERS-CoV spike protein, respectively. Therefore, it remains possible that the hedgehogs could harbor other MERS-like coronaviruses and transmit the virus to different species. The Merbecovirus genome sequences in humans possess around 65%-80% identity on average with other species, which also attributes to the efficient transmission, high diversity, and the significant amount of host species in Merbecovirus (Wong et al., 2019).

**Figure 2.**
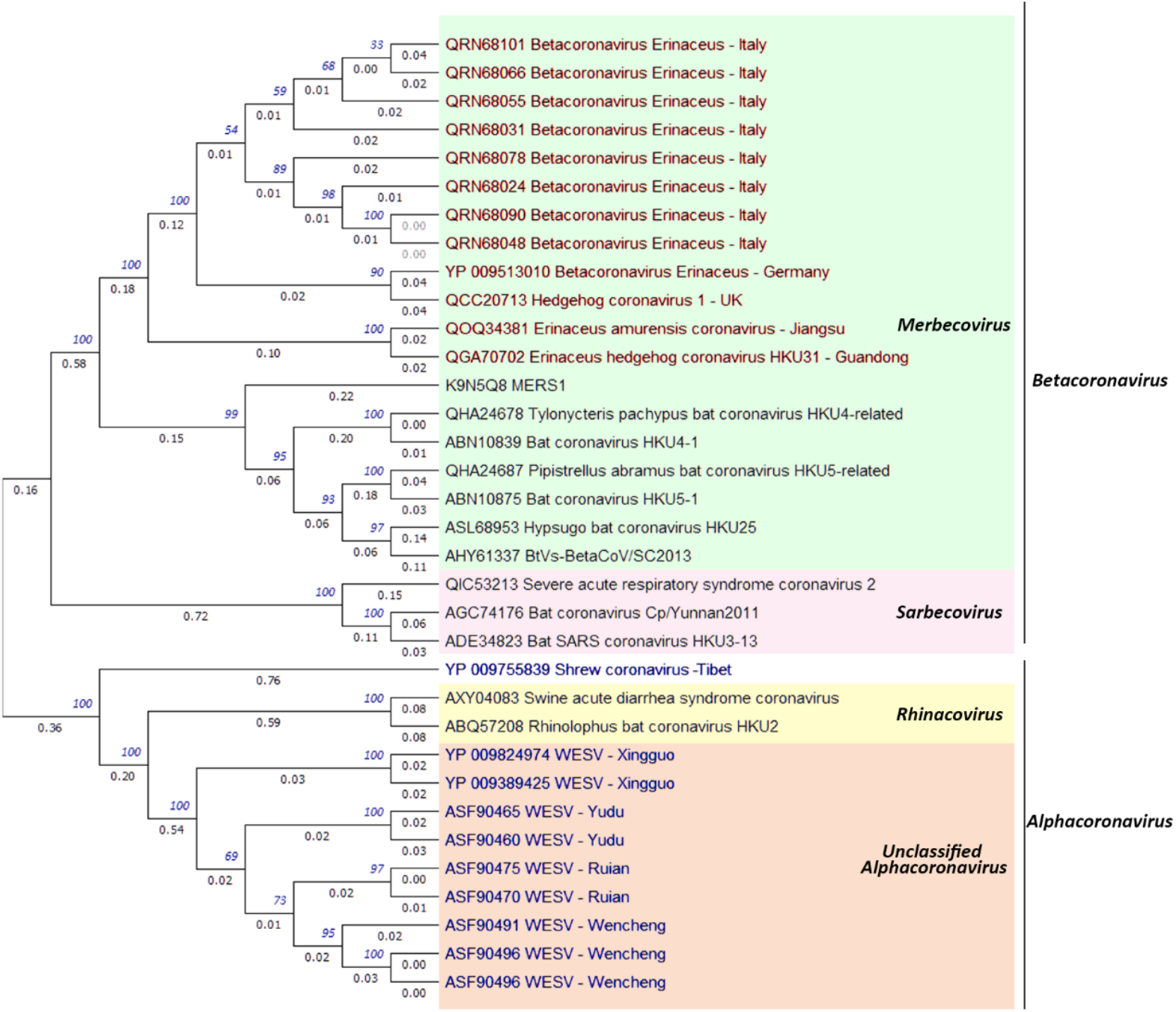
Maximum likelihood phylogenetic analysis of the hedgehog coronavirus stains, shrew coronavirus strains, and related alphacoronavirus and betacoronavirus strains based on spike protein amino acid sequences. The isolates colored in red are hedgehog coronaviruses and the isolates colored in blue are shrew coronaviruses

The amino acid sequence identities within WESVs are ranged from 90.05% to 99.74% through multiple sequence alignments. The group of WESV isolates forms a highly divergent group in the genus *Alphacoronavirus* based on the spike gene. The highest identity estimated between WESVs and other alphacoronaviruses is 34.68% with Rhinolophus bat coronavirus HKU2 from Rhinacovirus. Shrew-CoV/Tibet2014 itself, currently under unclassified Orthocoronavirinae, is a single clade under alphacoronavirus but distinct from WESV with only 25% identity of spike amino acid sequence. Its identity with other alphacoronaviruses is around 20% on average, which indicates its separate evolution in the genus.

Bats have been considered ancestors of alphacoronaviruses and betacoronaviruses by many studies (Drexler et al., 2014; Huynh et al., 2012; Vijaykrishna et al., 2007; Woo et al., 2012). However, WESVs and Shrew-CoV/Tibet2014 express highly divergent spike proteins against other alphacoronaviruses. Therefore, the evolution and ancestry of alphacoronaviruses might be more complicated than the results from former studies. Furthermore, the discovery of the novel coronavirus, Lucheng Rn rat coronavirus, also indicates the complexity of the evolutionary history of alphacoronaviruses (Wang et al., 2015).

### Geographical Distribution of the Coronaviruses from the Mammalian Order Eulipotyphla

As the phylogenetic analysis illustrates, the WESV isolates are clustered based on their geographic origins, which indicates the potential of in situ evolution of coronaviruses in Asian house shrews (Figure 3). The Western European hedgehog coronaviruses are centered in western Europe because of their host distributions. The Eulipotyphla coronaviruses have only been discovered in western Europe and China for now, which might be attributed to these animal species distributions and the limited research. Importantly, Europe, mainland Southeast Asia, Central Africa, as well as Central and South America, have been considered regions with high biodiversity of zoonotic infectious disease mammalian host species (Han et al., 2016). More studies should focus on these small mammals to shed light on the wildlife coronavirus evolution and zoonotic potential.

**Figure 3.**
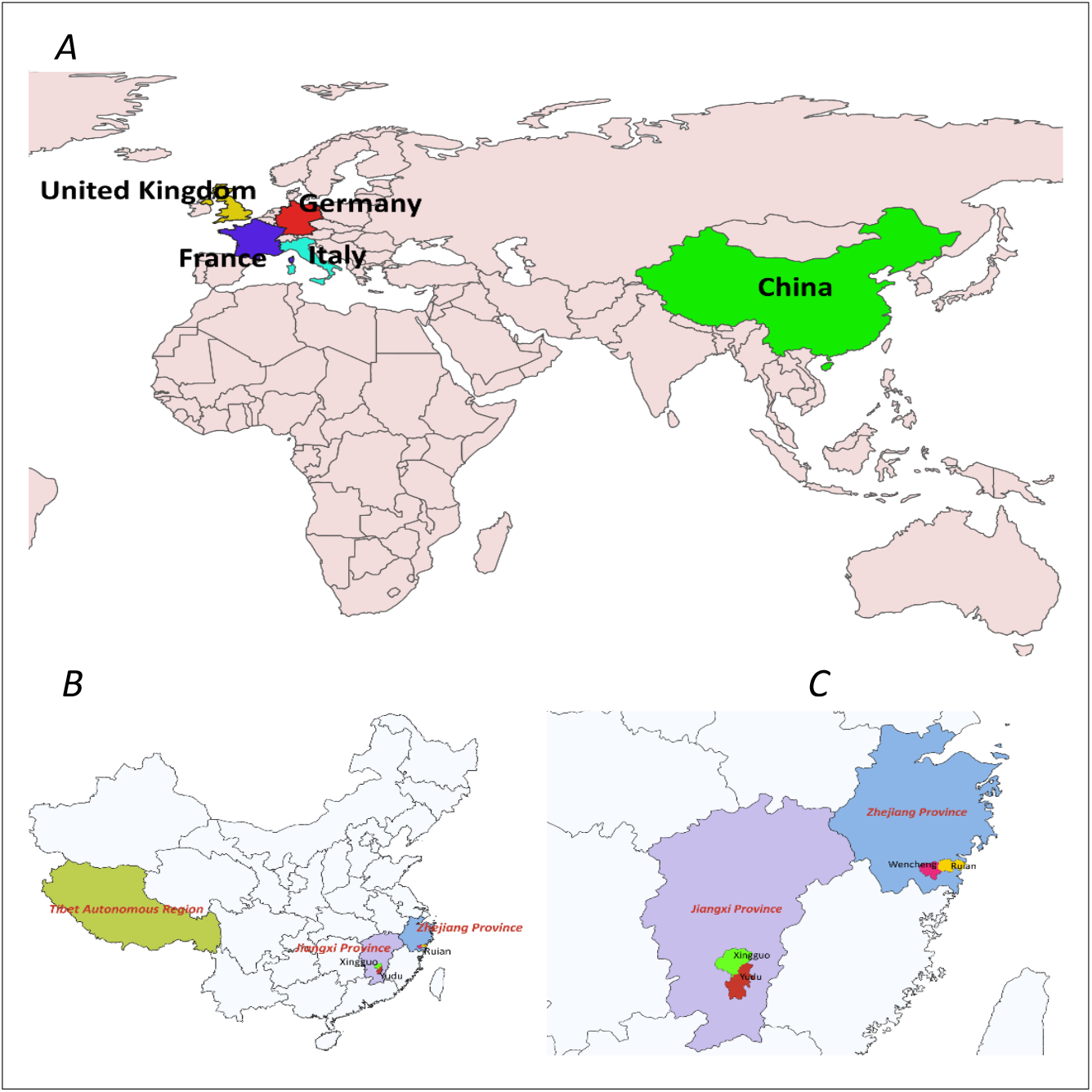
(A) Geographical distribution of the hedgehog and shrew (the mammalian order *Eulipotyphla*) coronavirus. The coronavirus strains are classified by countries: China (Wencheng Sm shrew coronavirus, Shrew coronavirus, Erinaceus hedgehog coronavirus HKU31, Erinaceus amurensis coronavirus); Germany (Betacoronavirus Erinaceus/VMC/DEU/2012); France (Erinaceus europaeus Alphacoronavirus); Italy (Betacoronavirus Erinaceus); UK (Betacoronavirus Erinaceus). (B) Geographical distribution of shrew coronaviruses. The shrew coronavirus strains are classified by provinces or regions: Tibet Autonomous Region (Shrew coronavirus); Zhejiang Province (Wencheng Sm shrew coronavirus); Jiangxi Province (Wencheng Sm shrew coronavirus). (C) Geographical distribution of Wencheng Sm shrew coronavirus. Wencheng Sm shrew coronavirus isolates are classified by counties: Yudu County (Yudu-76, Yudu-19); Xingguo County (Xingguo-101, Xingguo-74); Ruian County (Ruian-90, Ruian-133); Wencheng County (Wencheng-554, Wencheng-578, Wencheng-133).

### Furin cleavage Prediction

As furin cleavage is often assocated with spike protein activation and may be a key factor in zoonotic spillover and virus transmission, we analyzed 201 alphacoronavirus and betacoronavirus strains from the NCBI protein database for furin cleavage sites present on their spike proteins. Cleavage sites are found in 44 of them, with 17 hosted by bats and 27 by other mammals. The other 157 strains do not contain furin cleavage sites, with 143 hosted by bats and 14 by other mammals. The host animals and the number of coronaviruses they harbor are shown in Figure 4. The pie chart also includes some domestic animals that people have a big chance to contact, such as cats, dogs, pigs, horses, cattle, and rabbits. Therefore, we should pay attention to these animals as they have the potential to host transmissible coronaviruses.

**Figure 4.**
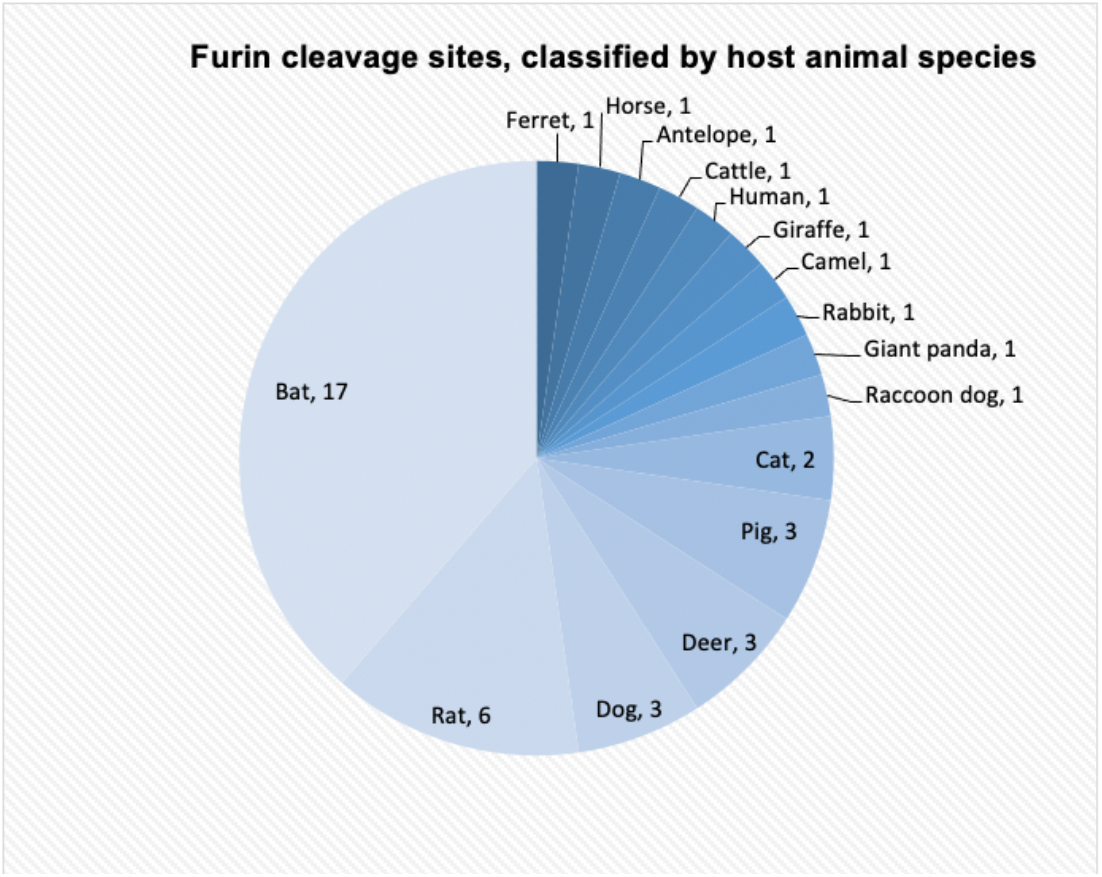
The number of alphacoronaviruses and betacoronaviruses with furin cleavage sites corresponding to their host animals. Most viruses are hosted by bats (17 strains).

We also examined the sequence of furin itself, as this can vary across animal species. Table 1 shows the pairwise sequence alignment result for furins across different animal species to see if there is any difference. The animals involved in the study are shrew, hedgehog, and the mammals shown in figure 4. Dolphin and greater horseshoe bat, whose furin proteases do not show any evidence to cleave the spike protein based on the surveillance study, are also included. The results show a high degree of conservation of furins across different animal species, which aligns with the previous study (El Najjar et al., 2015). The identities lower than 90% are between the camel and greater horseshoe bat, shrew and greater horseshoe bat, camel and horseshoe bat, and camel and hedgehog. Furins are ubiquitously expressed in various sites in all cells and tissues in eukaryotes, but little is known about whether the difference in furins across different species might affect their ability or efficiency to cleave the coronavirus spike protein (Hoffmann et al., 2018). Hence, more animals should be included, and more experiments need to be carried out to draw the conclusions (El Najjar et al., 2015; Huang et al., 2018).

**Table 1.** Pairwise sequence alignment result of furin across different animal species represented by percentage. Values in red are identities higher than 98%. Values in green are identities lower than 90%.

|  | human | camel | horse | dolphin | giant panda | ferret | cat | pig | cattle | greater horseshoe bat | fruit bat | rabbit | shrew | hedgehog |
| --- | --- | --- | --- | --- | --- | --- | --- | --- | --- | --- | --- | --- | --- | --- |
| human |  | 92.8 | 96.6 | 96.2 | 95.7 | 95.6 | 96.5 | 94.7 | 95.1 | 91.4 | 95.1 | 95.0 | 93.5 | 92.7 |
| camel | 92.8 |  | 93.0 | 92.6 | 91.7 | 91.4 | 92.0 | 91.4 | 92.4 | 87.6 | 91.7 | 90.5 | 90.4 | 88.7 |
| horse | 96.6 | 93.0 |  | 96.7 | 96.0 | 95.7 | 96.5 | 95.1 | 95.2 | 91.0 | 95.3 | 94.8 | 94.3 | 93.2 |
| dolphin | 96.2 | 92.6 | 96.7 |  | 96.0 | 95.4 | 96.5 | 95.5 | 95.6 | 90.8 | 94.6 | 94.2 | 93.7 | 93.2 |
| giant panda | 95.7 | 91.7 | 96.0 | 96.0 |  | 98.2 | 97.5 | 94.5 | 94.5 | 90.8 | 94.1 | 94.3 | 93.3 | 92.3 |
| ferret | 95.6 | 91.4 | 95.7 | 95.4 | 98.2 |  | 97.0 | 94.5 | 94.6 | 90.8 | 94.7 | 94.4 | 93.1 | 92.3 |
| cat | 96.5 | 92.0 | 96.5 | 96.5 | 97.5 | 97.0 |  | 95.0 | 95.2 | 91.2 | 94.7 | 94.7 | 93.6 | 93.5 |
| pig | 94.7 | 91.4 | 95.1 | 95.5 | 94.5 | 94.5 | 95.0 |  | 95.5 | 90.2 | 94.0 | 93.6 | 93.2 | 92.7 |
| cattle | 95.1 | 92.4 | 95.2 | 95.6 | 94.5 | 94.6 | 95.2 | 95.5 |  | 90.7 | 94.9 | 93.7 | 93.2 | 92.6 |
| greater horseshoe bat | 91.4 | 87.6 | 91.0 | 90.8 | 90.8 | 90.8 | 91.2 | 90.2 | 90.7 |  | 91.9 | 90.7 | 89.7 | 89.4 |
| fruit bat | 95.1 | 91.7 | 95.3 | 94.6 | 94.1 | 94.7 | 94.7 | 94.0 | 94.9 | 91.9 |  | 93.7 | 93.5 | 92.5 |
| rabbit | 95.0 | 90.5 | 94.8 | 94.2 | 94.3 | 94.4 | 94.7 | 93.6 | 93.7 | 90.7 | 93.7 |  | 92.9 | 92.5 |
| shrew | 93.5 | 90.4 | 94.3 | 93.7 | 93.3 | 93.1 | 93.6 | 93.2 | 93.2 | 89.7 | 93.5 | 92.9 |  | 92.1 |
| hedgehog | 92.7 | 88.7 | 93.2 | 93.2 | 92.3 | 92.3 | 93.5 | 92.7 | 92.6 | 89.4 | 92.5 | 92.5 | 92.1 |  |

### Prediction of Cleavage Sites in Hedgehog and Shrew Coronavirus Spike Proteins

With the presence of the furin cleavage sites, the fusion activation of the coronavirus is likely to be enhanced with increased infectivity, which broadens the host tropism range and enables the virus to be more transmissible. We estimated the furin cleavage sites of all Eulipotyphla coronaviruses by two programs, ProP and PiTou. The results run by ProP are displayed in Table 2. However, the analysis from PiTou (Tian et al., 2012) indicated that none of the viral strains have furin cleavage sites (not shown). Considering the different algorithms between these two programs, we followed up the results by ProP, based on predicted cleavage for furin family members. The general PC prediction stands for the general proprotein convertase cleavage sites prediction. The typical furin cleavage sites occur around S1/S2 (containing multiple arginines around site 700, such as **R**XX**R**) and S2’ (containing one or two arginines and the second **R** would be essential around 890) (Hoffmann et al., 2020; Kleine-Weber et al., 2018). For WESVs, six of them have positive ProP prediction results around site 513 and site 1121, which are not the typical S1/S2 or S2’ sites, so these sites might be cleaved by other proteases or be false positives. Yudu-76, Yudu-19, Ruian-90, Ruian-133, Xingguo-101, and Xingguo-74 isolates have a common potential cleavage site (**KR**|**S** around site 512) cleaved by proprotein convertases based on the general PC protection. The high scores in Yudu and Xingguo isolates suggest a high potential for the presence of these cleavage sites. They are likely to be cleaved by furin, but at atypical sites or by other furin-like proprotein convertases (PCs). Around 490 potential furin cleavage protein candidates were identified in the human proteome (Shiryaev et al., 2013). The PC family contains 7 members in human, comprising furin, PC1/3, PC2, PC4, PACE4, PC5/6, and PC7 (Remacle et al., 2008). PC5, PC7, and PACE4 are considered furin-like PCs. PACE4 is less likely to be the candidate for WESV cleavage sites because it cleaves at **KR** motifs less frequently than **R**X**K**/**RR** or **R**X**RR** (Gordon et al., 1997). PC5 has been demonstrated to cleave the SARS-CoV spike protein efficiently and it can cleave at **KR motifs**, indicating its potential to cleave WESV spike protein at **KR**|**S** (Basak et al., 2007). Though no studies have displayed the PC7 ability in cleaving coronavirus spike protein, it can cleave the unusual site **KR**|**S** (Corman et al., 2014). The endoproteases, cathepsin L and TMPRSS2, which can activate the SARS-CoV, MERS-CoV, and SARS-CoV-2 spike proteins, also have a likelihood to activate the WESV spike protein (Hoffmann et al., 2018; Matsuyama et al., 2020; Sacco et al., 2020). The studies on atypical furin cleavage sites and the corresponding PCs to specific viral proteins are still limited. Therefore, the spike protein model for WESV was further constructed to examine their furin cleavage site possibilities.

**Table 2.**
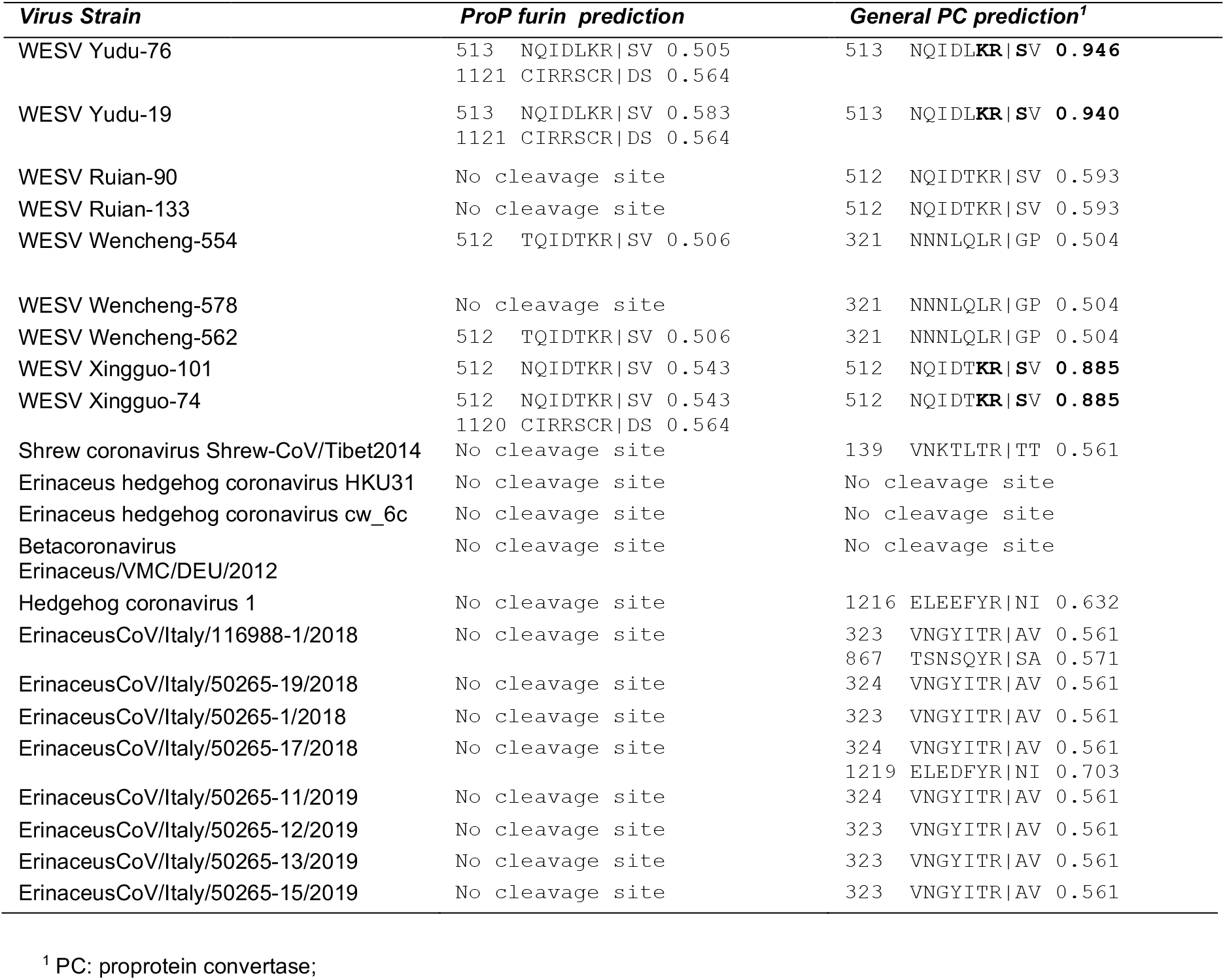
Prediction of cleavage sites in hedgehog and shrew coronavirus spike proteins. basic amino acids corresponding to positive scores above threshold (> 0.6) are shown in bold text

### Spike Protein Multiple Sequence Alignment

To better define the position of the putative cleavage sites, two representative WESV spikes (Ruian-90 and Yudu-76) showing high PC consensus sequence scores were aligned using MUSCLE alignment, and compared to MERS-CoV as a representative zoonotic coronavirus (see Appendix). Based on this, it is apparent that the putative PC cleavage sites do not align with the expected S1/S2 spike cleavage site, but rather occur in an upstream location within S1—being positioned broadly in line with a cleavage position previously identified in an unrelated rodent coronavirus, AcCoV-JC34 (discussed in Choi et al, 2021). As expected the conserved S2’ cleavage site was adjacent to the fusion peptide.

### Spike Protein Model Construction

To further examine the putative PC cleavage sites identified based on amino acid sequences, a WESV Yudu 76 spike protein structural model was constructed based on the HKU2 spike, which shares relatively high identity with WESVs (Figure 5). The cleavage site depicted in the structural model is clearly an exposed loop, which indicates it is in an appropriate environment to be accessible by proteases. One limitation of our work is that the amino acid identity of the reference model (*Rhinolophus* ba coronavirus HKU2) with WESV is relatively low (36%); however, we consider this to be a reliable model to illustrate the presence of a cleavage site—which, while it does not represent a typical furin cleavage site, may be cleaved by other furin-like proteases or trysin-like proteases. Further biochemical and in vivo assays are needed to determine the presence of functional cleavage sites for these viruses.

**Figure 5.**
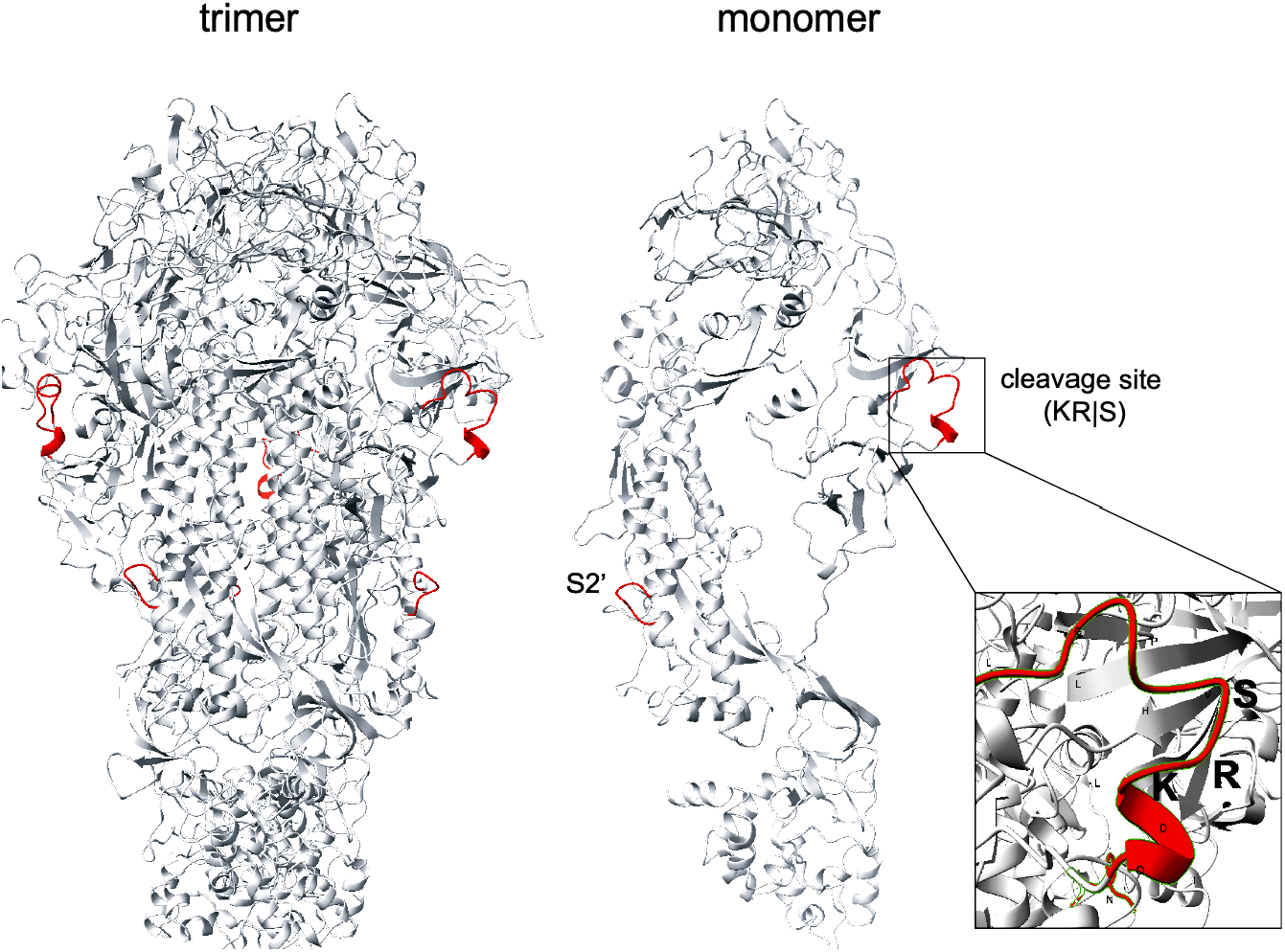
Wencheng Sm shrew coronavirus spike protein structure model, as a trimer and as a monomer. The model was constructed based on HKU2 spike glycoprotein (6M15). The predicted cleavage loop (QIDLKRSVHPVGL) is shown in red, with the proposed KR|S cleavage site indicated in the inset. The S2’ site and cleavage loop is also indicated.

### One Health Perspectives to Understand the Zoonotic Potential of Eulipotyphla Coronaviruses

Increasing zoonotic infectious diseases are affecting humans, domestic animals, and wildlife, in which case One Health approaches should be considered to reduce the risk of infectious diseases (Cunningham et al., 2017; Daszak et al., 2000). Here we integrated the factors involving the environment, wild animals, domestic animals, humans, and pathogens to discuss their interactions and the corresponding strategies to prevent the emerging coronaviruses.

Shrews are widely distributed in Asia, Africa, and Europe, sharing the same habitats with other insectivorous mammals and rodents (Bown et al., 2011). They tend to live in the rural and forested areas as well as the urban sewer drains (Rahman et al., 2018). Western European hedgehogs are endemic to the UK and Western Europe and mostly distributed in the gardens, woodlands, and grasslands (Hof et al., 2019). Amur hedgehogs have been found in China, Korea, and Russia so far (Ai et al., 2018; Lau et al., 2019). Their habitats include woodlands, grasslands, and also forest edges (Smith et al., 2010). Though many of these small mammals live in wild areas, the number of people who raise exotic companion mammals as pets is increasing all over the world (Rahman et al., 2018). Therefore, people need to be aware of the potential risk that exotic companion mammals like hedgehogs might carry a range of viruses when adopting them (Keeble & Koterwas, 2020).

The prevalence of coronaviruses in wildlife is not very high, due to the low density of animals caused by the dilution effect and restricted interspecies transmissions (Johnson & Thieltges, 2010; Zappulli et al., 2020). Wild animals require a certain receptor to harbor the virus, limiting the spread of the virus (Ji et al., 2020). The majority of the coronavirus wildlife host populations are rodents, insectivores like bats, and other mammals such as raccoon dogs, camels, pangolins, etc. (Irving et al., 2021). Typically, there are two wildlife-human interaction pathways. One pathway is that wild small mammals such as rodents might leave feces on crops, and farmers might get infected while handling these contaminated crops; the other is through the wildlife food supply trade, which is more prevalent (Huong et al., 2020). The initial SARS-CoV-2 transmissions were closely related to Wuhan wet food market (Tiwari et al., 2020). Various kinds of small mammals were sold either directly to the customers or to the restaurants, including hedgehogs, rodents, bats, pangolins and ferrets, which are all highly potential intermediate coronavirus hosts (Mahdy et al., 2020). Although shrews have not been demonstrated to be sold in the wet market, evidence has showed their presence with the rodents in the wet food market because of the poor sanitation and regulation (Tung et al., 2013).

A study on the viral infection persistence in temperate bats during their hibernation has shown that the coronavirus can persist for up to four months in laboratory setting (Subudhi et al., 2017). The reason might be that the bats conserve their energy and enter torpor with a low metabolic rate as well as low levels of inflammation during hibernation (Calisher et al., 2006; Subudhi et al., 2017). However, whether the wild bats have the equivalent ability of viral shedding is still not fully understood (Plowright et al., 2015). Hedgehogs also have the habits of hibernation during winter and share the nocturnal habits and insectivorous diets with bats (South et al., 2020). Therefore, we can speculate a high risk in hedgehogs to harbor and maintain the coronavirus during their hibernation, which raises the probability to transmit and circulate the virus among their living populations. Although there is no evidence that hedgehog and shrew coronaviruses are linked to any diseases, these mammals are considered hosts and potential natural reservoirs for coronaviruses. They might hold an important position in the virus evolution and have the ability to host newly emerging coronaviruses due to recombination and mutation of the coronaviruses.

Human activities, such as deforestation and conversion from land to agricultural use, drive the wild animals to move from rural areas to urban areas (McMahon et al., 2018). Climate change might also increase the insectivore populations feeding on insects due to the expanded insects (Morueta-Holme et al., 2010; Vega & Castro, 2019). More and more zoonotic epidemics occur due to these ecosystem changes, including the most recent SARS-CoV-2 and Ebola (Cunningham et al., 2017).

As a consequence of urbanization, humans are closer to wild animals than ever. For instance, Western European hedgehogs have been mostly found in backyards, urban grasslands, and woodlands (Hof et al., 2019). The interaction between domestic animals and wild animals is also becoming more frequent. For example, cats (the predators of the rodents and shrews), are likely to carry these wildlife coronaviruses during predation (Tsoleridis et al., 2016). The risk of zoonotic spillover is also elevated, leading to the spread of zoonotic epidemics (Hassani & Khan, 2020; Wang & Anderson, 2019). In Figure 6, an epidemiology triad was constructed to describe the process of how wildlife animal coronaviruses jump to humans due to urbanization.

**Figure 6.**
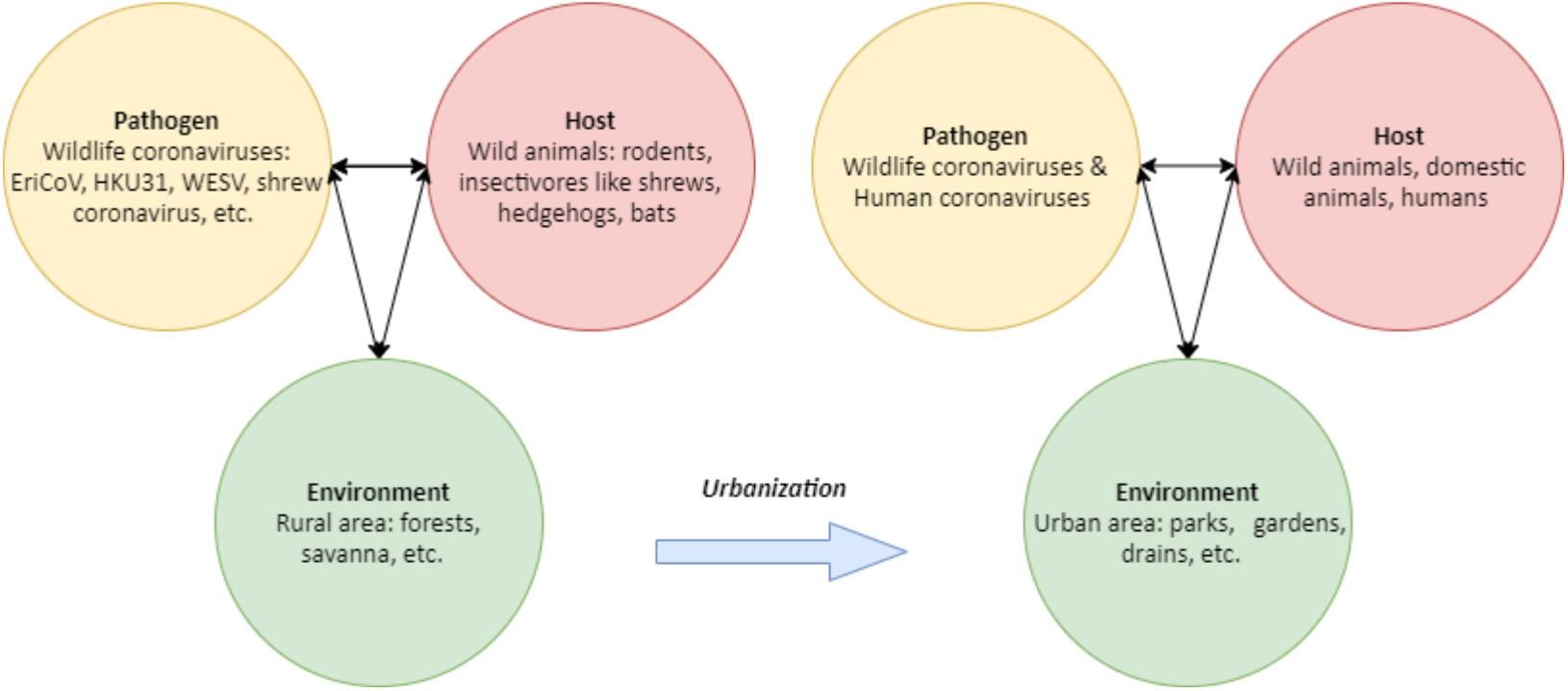
Epidemiology triad model before and after urbanization.

## Conclusions and Future Directions

This study conducted a bioinformatic analysis of the coronaviruses from the mammalian order Eulipotyphla, including hedgehogs and shrews. Shrew coronavirus and WESVs might diverge from a different ancestry compared to other alphacoronaviruses, revealing a more complicated alphacoronavirus evolution history than was previously thought. The hedgehog coronaviruses EriCoV and HKU31 are within the same subgenus Merbecovirus as MERS-CoV. WESV shows a PC cleavage site at **KR**|**S** around 512 site, indicating its zoonotic potential. The potential cleavage proteases could be PC5, PC7, and endoproteases other than furin. The animals, whose furins were demonstrated to have the ability to cleave the spike protein, are at a higher risk to host and transmit the coronaviruses, including bats, rats, dogs, deer, pig, cats, raccoon dogs, giant pandas, rabbits, camels, giraffes, cattle, horses, antelopes, and ferrets. No significance was found within the furins from a wide range of animals, but more studies are needed to demonstrate furin cleavage abilities. In conclusion, hedgehogs and shrews are natural reservoirs for coronaviruses, which might be able to recombine with other wildlife coronaviruses and generate new emerging viruses. As novel coronaviruses causes a heavy burden to global health, One Health perspectives and approaches are necessary to understand the evolution and transmission of the these viruses. Growing urbanization increases the wildlife to human interactions, and the small mammals sold in wet market mights also bring the pathogen spillover risks. Therefore, people should be vigilant of small wild mammals, and take precautions to control and prevent contact with these animals.

## ACKNOWLEDGEMENTS

We thank Michael Stanhope for invaluable help and advice in generating and analysing the data presented in this manuscript. A.C. is supported by grant T32EB023860 from the National Institute of Biomedical Imaging and Bioengineering. Work on coronavirus entry in the Whittaker lab is funded in part by the National Institute of Health research grant R01AI35270.

## Appendix

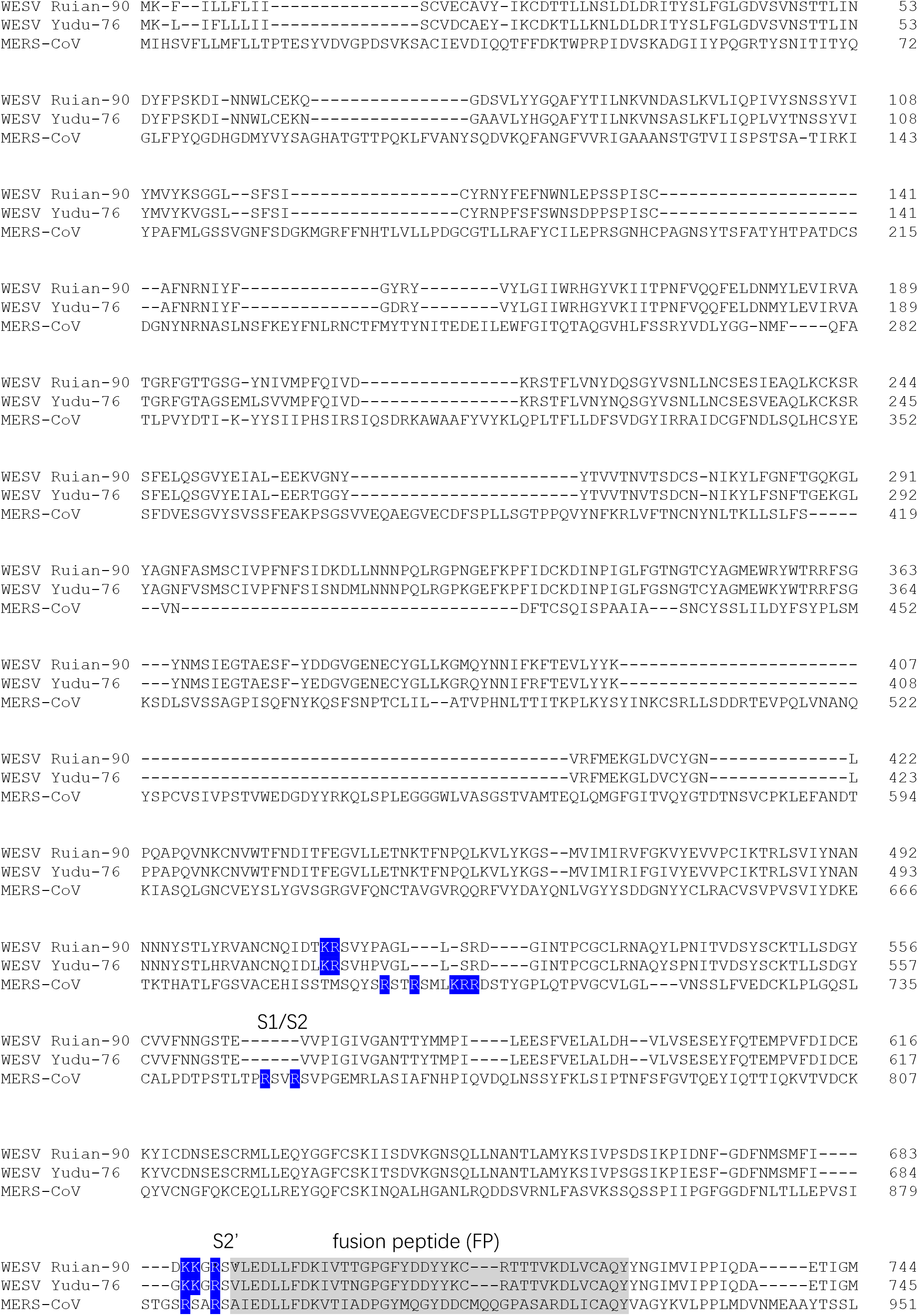
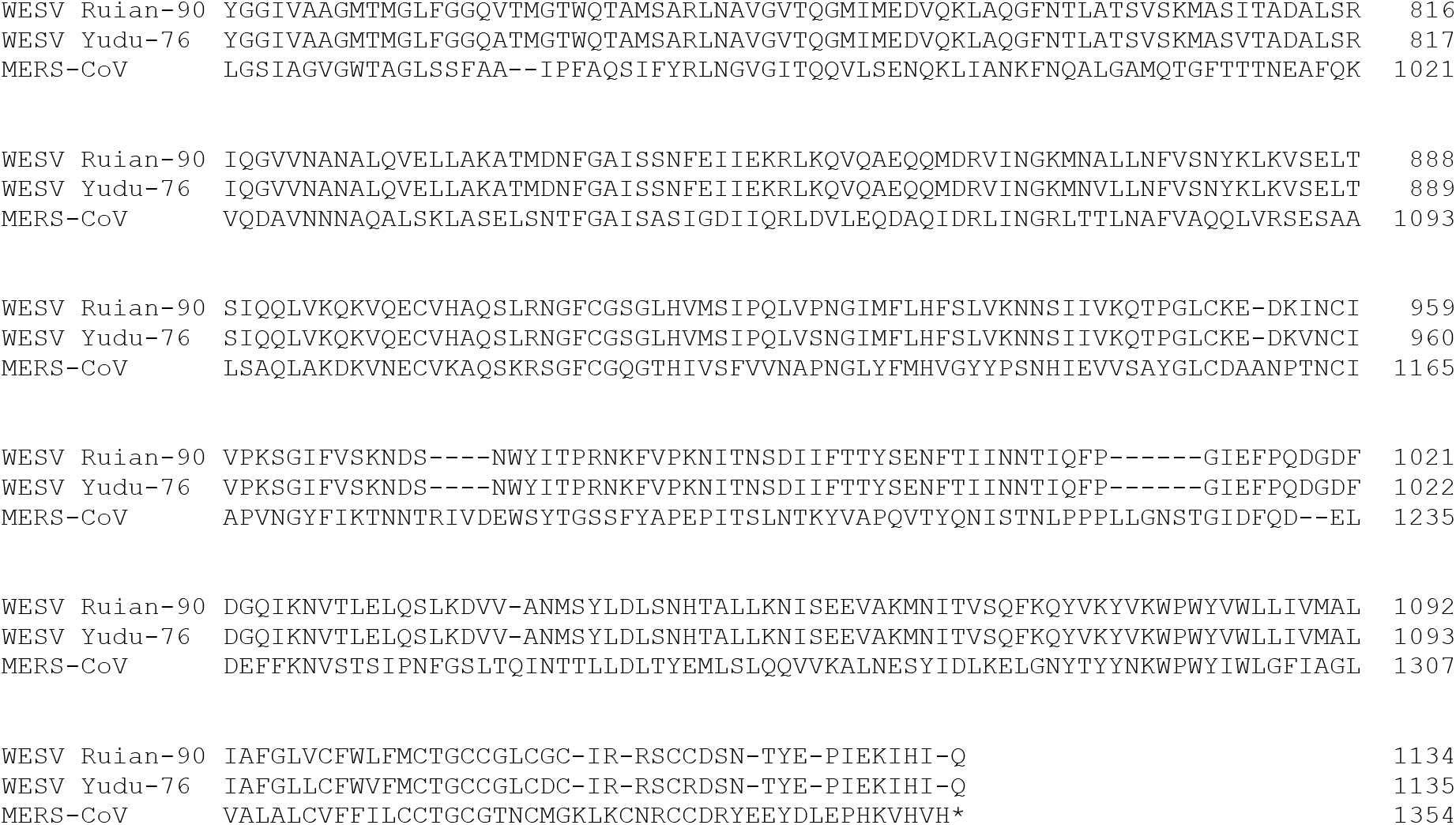

